# Glycolytic interference blocks influenza A virus propagation by impairing viral polymerase-driven synthesis of genomic vRNA

**DOI:** 10.1101/2022.11.09.515749

**Authors:** J. Kleinehr, K. Daniel, F. Günl, J. Janowski, L. Brunotte, M. Liebmann, M. Behrens, A. Gerdemann, L. Klotz, M. Esselen, H.-U. Humpf, S. Ludwig, E. R. Hrincius

**Affiliations:** Institute of Virology Muenster (IVM), Westfaelische Wilhelms-University Muenster, Von-Esmarch-Strasse 56, 48149 Muenster, Germany; Department of Neurology with Institute of Translational Neurology, University Hospital Muenster, Albert-Schweitzer-Campus 1, 48149 Muenster, Germany; Institute of Food Chemistry, Westfaelische Wilhelms-University Muenster, Corrensstrasse 45, 48149 Muenster, Germany

## Abstract

Influenza A virus (IAV), like any other virus, provokes considerable modifications of its host cell’s metabolism. This includes a substantial increase in the uptake as well as the metabolization of glucose. Although it is known for quite some time that suppression of glucose metabolism restricts virus replication, the exact molecular impact on the viral life cycle remained enigmatic so far. Using 2-deoxy-D-glucose (2-DG) we examined how well inhibition of glycolysis is tolerated by host cells and which step of the IAV life cycle is affected. We observed that effects induced by 2-DG are reversible and that cells can cope with relatively high concentrations of the inhibitor by compensating the loss of glycolytic activity by upregulating other metabolic pathways. Moreover, mass spectrometry data provided information on various metabolic modifications induced by either the virus or agents interfering with glycolysis. In the presence of 2-DG viral titers were significantly reduced in a dose-dependent manner. The supplementation of direct or indirect glycolysis metabolites led to a partial or almost complete reversion of the inhibitory effect of 2-DG on viral growth and demonstrated that indeed the inhibition of glycolysis and not of *N*-linked glycosylation was responsible for the observed phenotype. Importantly, we could show via conventional and strand-specific qPCR that the treatment with 2-DG led to a prolonged phase of viral mRNA synthesis while the accumulation of genomic vRNA was strongly reduced. At the same time, minigenome assays showed no signs of a general reduction of replicative capacity of the viral polymerase. Therefore, our data suggest that the significant reduction in IAV replication by glycolytic interference occurs mainly due to an impairment of the dynamic regulation of the viral polymerase which conveys the transition of the enzyme’s function from transcription to replication.

**Author Summary:** Upon infection the influenza A virus alters the metabolism of infected cells. Among others, this includes a pronounced increase in glucose metabolism. We aimed to get a better understanding of these metabolic virus-host interactions and to unravel the mechanism by which glycolytic inhibition impairs the viral life cycle. On the one hand, we observed a virus-induced upregulation of many glycolysis metabolites which could often be reversed by the administration of a glycolysis inhibitor. On the other hand, our data suggested that the inhibitor treatment severely impaired viral propagation by interfering with the regulation of the viral polymerase. This manifested in an extended phase of transcription, while replication was strongly reduced. Additionally, we assessed the safety and tolerability of the used drug in immortalized and primary cells. Our study sheds more light on metabolic virus-host interactions and provides a better understanding of metabolic interference as a potential host-targeted antiviral approach, which does not bear the risk of creating resistances.

## 1. Introduction

Influenza viruses (IVs) still constitute a major risk factor for the human health all over the globe. According to extrapolations, 3-5 million severe cases and up to half a million deaths occur on average during annual IV epidemics [1]. The influenza A virus (IAV) is of special interest since it has zoonotic and pandemic potential. The high mutation rate of the IV genome easily allows to develop resistances to antiviral treatments. Therefore, more and more research focuses on targeting cellular factors, which are indispensable for viral replication, to develop novel host-targeted antiviral strategies. Since viruses in general are intracellular parasites and thus have no metabolism on their own, they completely depend on the host cell’s metabolism for their replication. Moreover, each type of virus reshapes the host cell’s metabolism towards its specific needs by regulating – often increasing – the uptake of metabolites and the activity of certain metabolic pathways [2-6]. Frequently, this includes elevated activity of glycolysis, the pentose phosphate pathway (PPP), lipid metabolism and the generation of amino acids [3]. This was also demonstrated for IV infections. Altered activity or elevated levels of pathway intermediates of, among others, glutaminolysis [7-9], fatty acid synthesis (FAS) [7, 9], the PPP [7, 8], the hexosamine biosynthetic pathway [9] and the tricarboxylic acid (TCA) cycle [7, 8] were observed. However, especially an increased glycolytic rate and uptake of glucose has been described in various immortalized and primary cells after infection with IV as well as in the lungs of infected patients [7, 8, 10]. Direct inhibition of glycolysis or mediators of glycolysis led to a significant impairment of IV reproduction and spread [7, 11, 12]. Furthermore, the concentration of extracellular lactate increases during IV infections [8], suggesting the exploitation of aerobic glycolysis. This is indicative of the Warburg effect [13, 14], in which cells metabolize glucose rather to lactate instead of pyruvate despite the adequate availability of oxygen. In this scenario, which is also observed in tumors, cells depend more on glycolysis than oxidative phosphorylation for sufficient synthesis of adenosine triphosphate (ATP). On the one hand IV benefits from this by rapidly generating large amounts of biological building blocks for its replication and on the other hand an increased production of lactate inhibits the induction of type I interferons [15].

In our research we targeted the glucose metabolism with a special focus on the inhibition of glycolysis with the inhibitor 2-deoxy-D-glucose (2-DG), which has already been demonstrated to interfere with the formation of new infectious IV particles [11, 12, 16, 17]. Beside the competitive inhibition of glucose uptake, 2-DG inhibits the first two glycolytic enzymes hexokinase (HK) and glucose-6-phosphate isomerase (GPI), the latter being its primary target. Just like glucose, 2-DG will be phosphorylated at the C6 position by HK to 2-deoxy-D-glucose-6-phosphate (2-DG-6-P). 2-DG-6-P competitively inhibits GPI and cannot be further metabolized by this enzyme. The increasing concentration of 2-DG-6-P leads to a feedback that additionally inhibits hexokinase in an allosteric manner [18-22]. Moreover, 2-DG gets fraudulently incorporated into oligosaccharide chains needed for *N*-linked glycosylation of glycoproteins [23], partially preventing this post-translational modification [24] and hence affecting the proteins’ folding and their functions. This inhibition is mainly conveyed by guanosine diphosphate (GDP)-2-DG into which 2-DG can be converted [25]. Thereby, 2-DG evidentially inhibits glycolysis and interferes with *N*-linked glycosylation. Here, we demonstrate the inhibitor’s significant impact on the replication of IAV without causing irreversible damage to the host cells. Furthermore, we unraveled a major mechanism by which this treatment interferes with the viral life cycle and discuss the potential of metabolic interference to fight severe IAV infections.

## 2. Results

### 2-DG is well tolerated in cells and exhibits strong virus-restricting activity

Our first aim was to prove the virus-restricting potential of 2-DG in cell culture. First, we showed in plaque assays that the number of newly produced infectious IAV particles decreased significantly in a dose-dependent manner when 2-DG was applied directly after the infection of A549 cells (**Fig 1A**). This decrease became as strong as more than four orders of magnitude when the glucose/2-DG ratio was 1:1. Second, we observed a very similar 2-DG-mediated decrease for IAV nucleoprotein (NP)-positive cells via flow cytometry (**Fig S1A**). These data demonstrated the strong impairment of IAV reproduction and spread in the presence of 2-DG.

**Fig 1:**
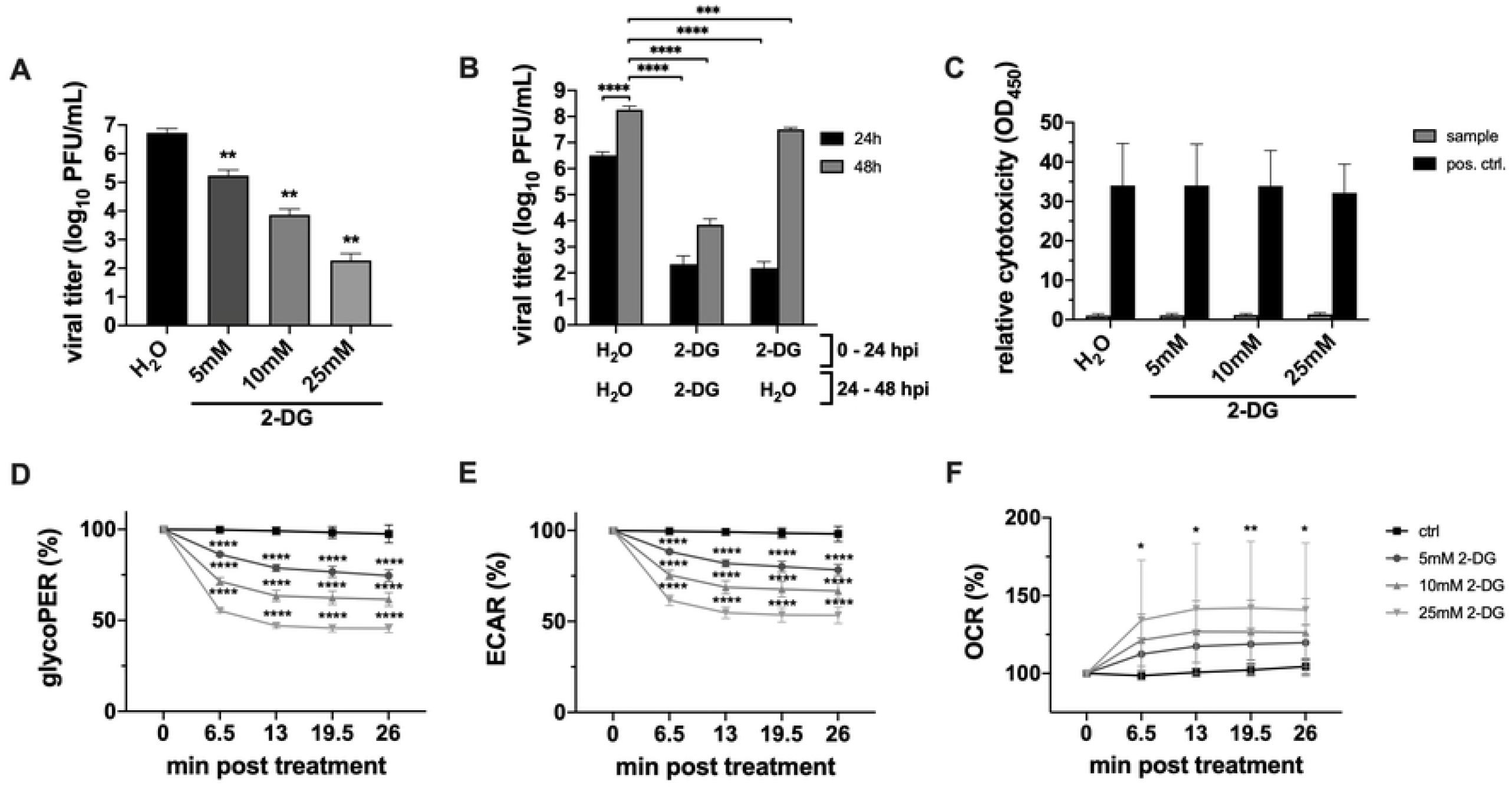
2-DG impairs IAV propagation and is well tolerated by A549 cells. **(A+B)** 24 h after seeding, A549 cells were infected with SC35M at an MOI of 0.001 for 30 min and incubated in the presence of 25 mM glucose and the indicated concentrations of 2-DG or its solvent water for **(A)** 24 h or **(B)** 24 and 48 h. Subsequently, supernatants were collected to determine viral titers via plaque assay. **(C)** Uninfected cells were treated with the indicated inhibitor concentrations for 24 h and were then subjected to LDH assay for assessment of the relative cytotoxicity of the treatment. **(D-F)** The glycolytic proton efflux rate (glycoPER), extracellular acidification rate (ECAR) and oxygen consumption rate (OCR) were measured in real-time via glycolytic rate assay and a Seahorse XFe96 Analyzer. The kinetics show the influence of different concentrations of 2-DG on the three measured parameters. Depicted are the means ± SD of **(A-C)** three or **(D-F)** five independent experiments with **(A-C)** three or **(D-F)** four biological replicates per condition and experiment. Statistical significances were determined via **(A)** unpaired one-way ANOVA and Dunnett’s correction, comparing all treated samples to the water control and **(B-F)** ordinary two-way ANOVA with **(B)** Tukey’s, **(C)** Sidak’s and **(D-F)** Dunnett’s correction for multiple comparison, comparing **(B)** all samples with one another, **(C)** all treated samples of one group to the respective water control or **(D-F)** the time points of differentially treated cells with their respective start value. p-values are indicated as follows: < 0.05 = *, < 0.01 = **, < 0.001 = ***, < 0.0001 = ****.

Next, we assessed the reversibility as well as metabolic and potential cytotoxic effects of the 2-DG treatment on cells. Here, it could be demonstrated that the strong antiviral effect of a 24 h treatment was quickly abolished once the inhibitor was removed (**Fig 1B**). The massive increase of viral titers after the replacement of 2-DG with inhibitor-free medium suggested the full reversibility of 2-DG-induced effects and indicated that there was no permanent cell damage which is also substantiated by the literature [11]. By performing lactate dehydrogenase (LDH) assays we detected no cytotoxicity within the range of used 2-DG concentrations (**Fig 1C**), as previously demonstrated in various cell lines including A549 [26, 27]. Moreover, we could even observe a beneficial effect of the 2-DG treatment for the survival of infected cells. With increasing 2-DG concentrations the total percentage of dead cells decreased significantly 24 hours post infection (hpi) (**Fig S1B**). However, the results of the LDH assays in combination with data obtained from trypan blue exclusions suggested a certain cytostatic effect, since even though the viability of all samples was not affected, total cell counts decreased with rising 2-DG concentrations (**Fig S1C+D**). In line with these results, a cytostatic effect of 2-DG has also been observed previously in other cells [27-29]. Furthermore, we investigated the effect of 2-DG on the metabolism in real-time via a Seahorse Analyzer. We observed a very rapid and significant reduction of the glycolytic proton efflux rate (glycoPER), which constitutes a direct read-out of the glycolytic rate (**Fig 1D**). Since a major factor to calculate the glycoPER is the extracellular acidification rate (ECAR), this decreased in a similar pattern as the glycoPER (**Fig 1E**). Simultaneously, the oxygen consumption rate (OCR) of 2-DG-treated cells increased quickly after the beginning of the treatment (**Fig 1F**). These data proved the partial inhibition of glycolysis by 2-DG in a dose-dependent manner and indicated that cells were able to cope with the treatment by compensating the loss of glycolytic activity by upregulating cellular respiration to generate energy.

In addition to evaluating the cytotoxicity of 2-DG, we also tested potential effects of 2-DG on the innate immune response and the cellular responsiveness to viral infections. For that purpose, we measured expression levels of the proinflammatory genes *interleukin-6* (*IL-6*) and *C-X-C motif chemokine ligand 8* (*CXCL8*, protein: IL-8) as well as the interferon-stimulated genes (ISGs) *DExD/H-box helicase 58* (*DDX58*, protein: retinoic acid inducible gene I) and *myxovirus resistance gene A* (*MxA*) after stimulation with either cellular or viral RNA in the presence or absence of 2-DG (**Fig S1E-H**). We observed a mild to more pronounced induction of *IL-6* (**Fig S1E**) and *CXCL8* (**Fig S1F**) with increasing concentrations of 2-DG. This finding was consistent with a previous publication, reporting that nutrient shortage (also induced by 2-DG) triggers a cell response which resembles wound healing processes in cancer cells as well as in primary cells [26]. Moreover, the mild induction of proinflammatory cytokines in the presence of 2-DG might be attributed to the fact that the inhibitor can also impair glycosylation. This in turn gives rise to endoplasmic reticulum (ER) stress, elicited by deficient glycoproteins, consequently leading to the unfolded protein response (UPR) [23] which has been demonstrated to drive the production of proinflammatory cytokines [30]. On the other hand, we measured no clear differences in the expression of *DDX58* (**Fig S1G**) and *MxA* (**Fig S1H**) in the presence of lower 2-DG concentrations but a moderate and significant reduction of both ISGs at 25 mM of the inhibitor, when stimulated with viral RNA. Nevertheless, our data confirmed that the cells were well responsive to viral stimuli, regardless of the concentration of 2-DG that was applied.

Apart from the permanent cell line A549, key experiments were repeated in primary human bronchial epithelial cells (HBEpCs) and genuine human lung explants (**Fig S2A-E**). Since the used media for primary tissue contained less glucose, lower concentrations of the inhibitor were used. However, we still applied the same 2-DG/glucose ratio to human lung explants as in A549 experiments which led to a significant and dose-dependent reduction of viral titers (**Fig S2A**). Because HBEpCs were more susceptible to the treatment, lower 2-DG/glucose ratios were applied. The highest concentration used in HBEpC experiments was 1200 μM which corresponds to the 2-DG/glucose ratio (1:5) of 5 mM 2-DG in experiments carried out with A549 cells. Similar to A549 cells, HBEpCs displayed barely any signs of cytotoxicity after treatment (**Fig S2B**). Reduced lactate concentrations in the supernatant of treated cells indirectly indicated the efficiency of glycolytic inhibition (**Fig S2C**). Importantly, the treatment with 2-DG also led to a significant and dose-dependent reduction of viral titers in HBEpCs (**Fig S2D+E**). Even though the magnitude of the inhibitory effect on glycolysis and viral replication differed slightly from the data obtained with A549 cells – most likely due to distinct cellular metabolic activities and lower 2-DG/glucose ratios (HBEpC) – these data suggested the save use and antiviral activity of 2-DG in primary tissue.

### 2-DG only moderately affects viral protein translation in a single viral life cycle

Given the remarkable impairment of IAV replication by 2-DG, we now aimed to identify the spot of interference of the drug within the viral life cycle. Therefore, we checked potential changes in the accumulation of various IAV proteins 24 hpi (**Fig 2A**) and after a single replication cycle of 8 h (**Fig 2B**). In accordance with the strongly reduced viral titers there was also a pronounced reduction of viral protein accumulation after 24 h. However, within a single replication cycle we only detected rather weak differences among the accumulation of viral proteins. While the accumulation of the late viral proteins polymerase acidic protein (PA) and matrix protein 1 (M1) was moderately reduced, the accumulation of the early proteins NP and non-structural protein 1 (NS1) was barely affected by 2-DG. Consequently, even though there was a moderate reduction of some viral proteins within a single replication cycle we did not consider reduced viral protein accumulation to be the main reason for the severe impact of 2-DG on IAV propagation.

**Fig 2:**
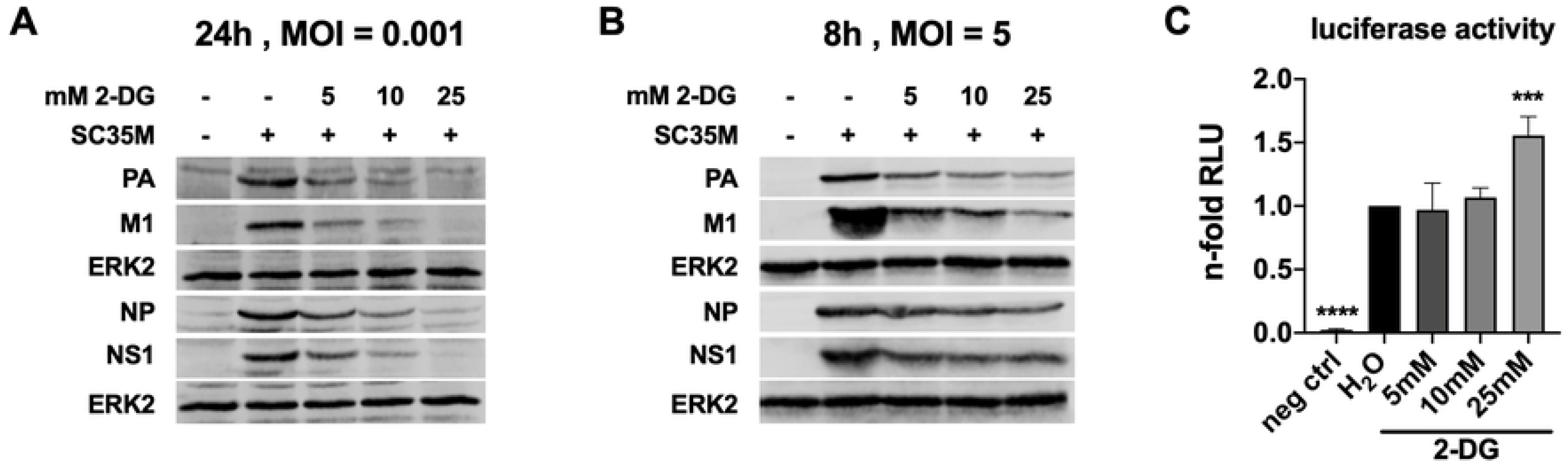
2-DG mildly reduces the expression of viral proteins. **(A+B)** 24 h after seeding, A549 cells were infected with SC35M at the depicted MOIs for 30 min and were incubated with 25 mM glucose and the indicated concentrations of 2-DG for a total of 24 h or **(B)** 8 h. Lysates of triplicates were unified to yield sufficient protein amounts. Proteins were separated via SDS-PAGE. Visualization was done using primary antibodies against PA (rabbit), M1 (mouse), NP (rabbit), NS1 (rabbit) and ERK2 (rabbit) and fluorescence-labelled anti-mouse (donkey) and anti-rabbit (donkey) secondary antibodies. Depicted are representative protein bands from three independent experiments. **(C)** HEK293T cells were, 24 h after seeding, transfected with an empty vector or a plasmid containing the *Renilla* luciferase gene which is under the control of a constitutive herpes simplex virus thymidine kinase promoter. Subsequently, the cells were incubated with the shown 2-DG concentrations. After 24 h, cells were lysed and the n-fold of relative light units (RLU) in comparison to the water control was measured via luciferase assay. Depicted are the means ± SD of three independent experiments with three biological replicates per condition and experiment. Statistical significances were determined via unpaired one-way ANOVA and Dunnett’s correction, comparing all treated samples to the water control. p-values are indicated as follows: < 0.05 = *, < 0.01 = **, < 0.001 = ***, < 0.0001 = ****.

To rule out a general effect on the cellular protein synthesis machinery, we measured the fluorescence signal of the reporter *Renilla* luciferase driven by a constitutive promoter in a luciferase assay in the absence or presence of various concentrations of 2-DG (**Fig 2C**). Decreased signals would be an indication for an impairment of cellular transcription and/or translation. Interestingly, there was no negative effect on the luciferase signal, suggesting no general impairment of the cellular protein synthesis. Quite the opposite was the case when high concentrations of 2-DG were used which even led to an increase of the luciferase signal.

### Glycolytic interference prolongs the phase of viral transcription while it clearly reduces viral replication within a replication cycle

After disproving viral protein expression being notably hampered by 2-DG, we delved deeper into the IAV replication cycle to understand the virus-restricting properties of 2-DG. Therefore, we now examined if a treatment with 2-DG interfered with the main processes driven by the viral polymerase: transcription and replication. Since IAV is a negative-sense RNA virus its RNA-dependent RNA polymerase can, right after reaching the host cell’s nucleus, transcribe positive-sense mRNA. After translation and nuclear import, nascent viral polymerase complexes mediate the two-step process of replication. Here, a positive-sense, full-length complementary RNA (cRNA) is synthesized from the initial viral genomic RNA (vRNA) which subsequently serves as a template for vRNA synthesis [31, 32].

We analyzed the accumulation of viral mRNA and vRNA that codes for M1. In case of vRNA detection, the values of M1 are representative of segment 7 (M). As before, M1 mRNA and vRNA were analyzed after 24 h (**Fig 3A+B**) and after a single replication cycle of 8 h (**Fig 3C+D**) with and without 2-DG. As observed for viral proteins, we measured a massive reduction of M1 mRNA and vRNA 24 hpi when 2-DG was applied (**Fig 3A+B**), which is in line with the reduction of viral titers. Experiments for the duration of a single replication cycle, however, revealed intriguing differences between the two distinct RNA species. While viral mRNA levels were elevated in the presence of 2-DG (**Fig 3C**) the amount of vRNA was clearly reduced after an infection period of 8 h (**Fig 3D**). Again, these experiments were repeated with HBEpCs to see if there are similar effects in non-transformed cells with no altered metabolism (**Fig S2F-I**). Using these primary cells, we observed a very similar pattern of IAV mRNA and vRNA accumulation through the treatment with 2-DG as in A549 cells. While mRNA was decreased 24 hpi (**Fig S2F**) and unaffected 8 hpi (**Fig S2G**), vRNA was decreased at both time points (**Fig S2H+I**). The difference in mRNA accumulation 8 hpi might be due to a milder 2-DG treatment or could be a cell-specific effect. Nevertheless, the strong reduction in vRNA accumulation, limiting viral propagation, seemed to be tissue-independent.

**Fig 3:**
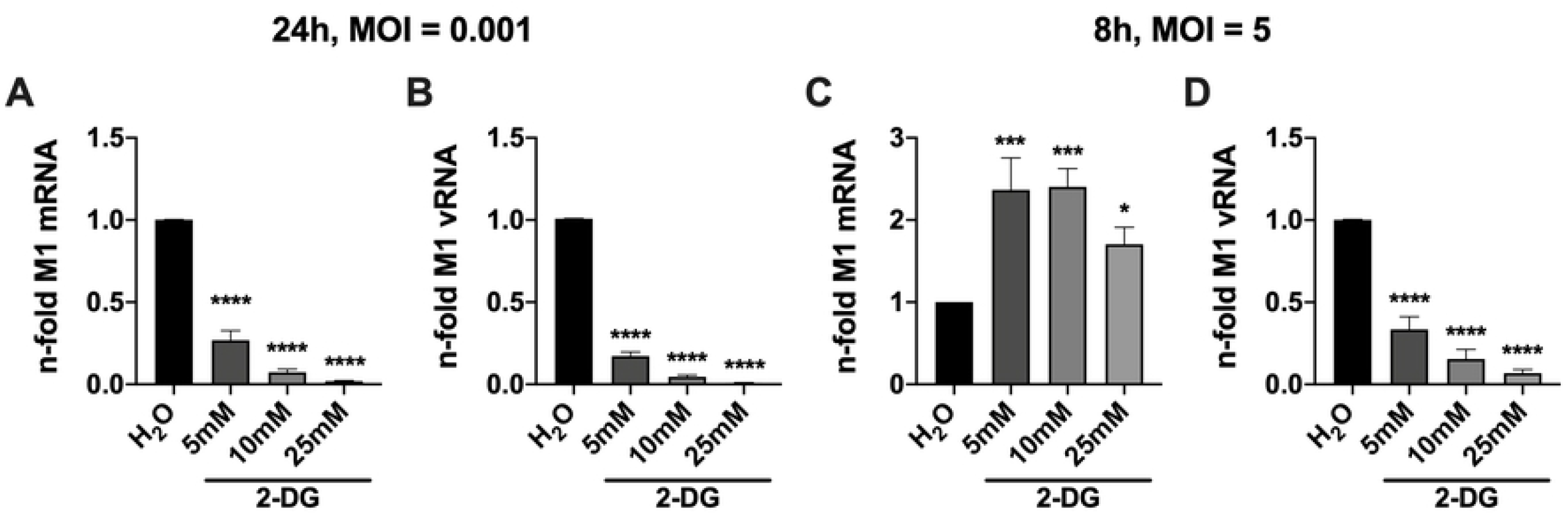
2-DG conversely affects IAV mRNA and vRNA accumulation. 24 h after seeding, A549 cells were infected with SC35M at the depicted MOIs for 30 min and were incubated with 25 mM glucose and the indicated concentrations of 2-DG for a total of **(A+B)** 24 h or **(C+D)** 8 h. Subsequently, cells were lysed, their RNA isolated and cDNA synthesized using either **(A+C)** oligo(dT) primers to transcribe mRNA or **(B+D)** fluA uni12 primers to transcribe vRNA. Real-time qPCR was performed with two technical replicates per sample and values of treated samples were normalized to the water control. In case of mRNA detection, all results were additionally normalized to a GAPDH control. Depicted are the means ± SD of three independent experiments with three biological replicates per condition and experiment. Statistical significances were determined via unpaired one-way ANOVA and Dunnett’s correction, comparing all treated samples to the water control. p-values are indicated as follows: < 0.05 = *, < 0.01 = **, < 0.001 = ***, < 0.0001 = ****.

With this phenotype at hand, we wanted to exclude a virus strain-specific effect and additionally analyzed the influence of 2-DG on viral growth, transcription, and replication of the H3N2 strain A/Panama/2007/1999 (Pan/99). As for SC35M, we observed a strong dose-dependent decrease of viral titers, mRNA and vRNA 24 hpi (**Fig S3A-C)**. Importantly, with an increase of Pan/99 mRNA and a decrease of vRNA in a single cycle experiment **(Fig S3D+E)** the results resembled those obtained with SC35M. Therefore, glycolytic interference on IAV appears to be a general phenomenon and not a virus strain-specific effect.

Summing up the obtained insights, the qPCR data suggested that the main cause for the impairment of IAV reproduction and spread by 2-DG is the interference of the inhibitor with the production of viral genome copies. Hereafter, we were especially interested in why glycolytic inhibition increased viral mRNA but decreased vRNA within a single viral life cycle.

In order to shed light on this question we performed an 8 h infection kinetic and analyzed the synthesis of M1 mRNA and vRNA in the presence of 2-DG in comparison to an untreated control (**Fig 4A+B**). In untreated cells the production of viral mRNA reached its strongest incline at approximately 6 hpi and started to establish a plateau afterwards (**Fig 4A**, black line). In contrast, the treatment with 2-DG led to a continuous increase of mRNA transcription, exceeding the total accumulation of viral mRNA in untreated cells (**Fig 4A**, gray line). Thus, despite a lower accumulation rate of viral mRNA in treated cells in the first 6 h of an infection, these samples displayed higher mRNA levels at time points later than 7 hpi. Even though the underlying mechanisms are unknown this observation explained why we detected higher viral mRNA levels in 2-DG-treated cells after one replication cycle (**Fig 3C**). In accordance with our previous data on vRNA accumulation at 8 hpi (**Fig 3D**), the kinetic revealed that vRNA accumulated at a clearly reduced rate when 2-DG was applied throughout the whole experiment (**Fig 4B**, gray line). To verify our results, we performed strand-specific real-time qPCR according to the protocol established by Kawakami *et al*. [33] for segment 5 (NP) and 6 (NA). Additionally, we analyzed segment 1 (PB2), which is the longest of the IAV gene segments, to rule out effects which might be caused by the length of different segments. We determined the n-fold of viral mRNA and vRNA of the three segments in 2-DG-treated cells 8 hpi in comparison to untreated cells. The results for all three gene segments were very similar and supported the previous kinetics. We observed a 3-4-fold increase of viral mRNA (**Fig 4C-E**) while the vRNA of the same gene segments was decreased by approximately 80-90% (**Fig 4F-H**) when 2-DG was applied. Notably, these findings confirmed our previous measurements of mRNA and vRNA after one replication cycle (**Fig 3C+D**). The data presented in **Fig 4** indicated that glycolytic inhibition by 2-DG prolonged the phase of viral mRNA transcription while it attenuated viral genome replication. This suggested either a distinct effect on the transcriptional and replicative capacity of the viral polymerase or an impairment of the dynamic regulation of the polymerase function, determining whether it performs transcription or replication.

**Fig 4:**
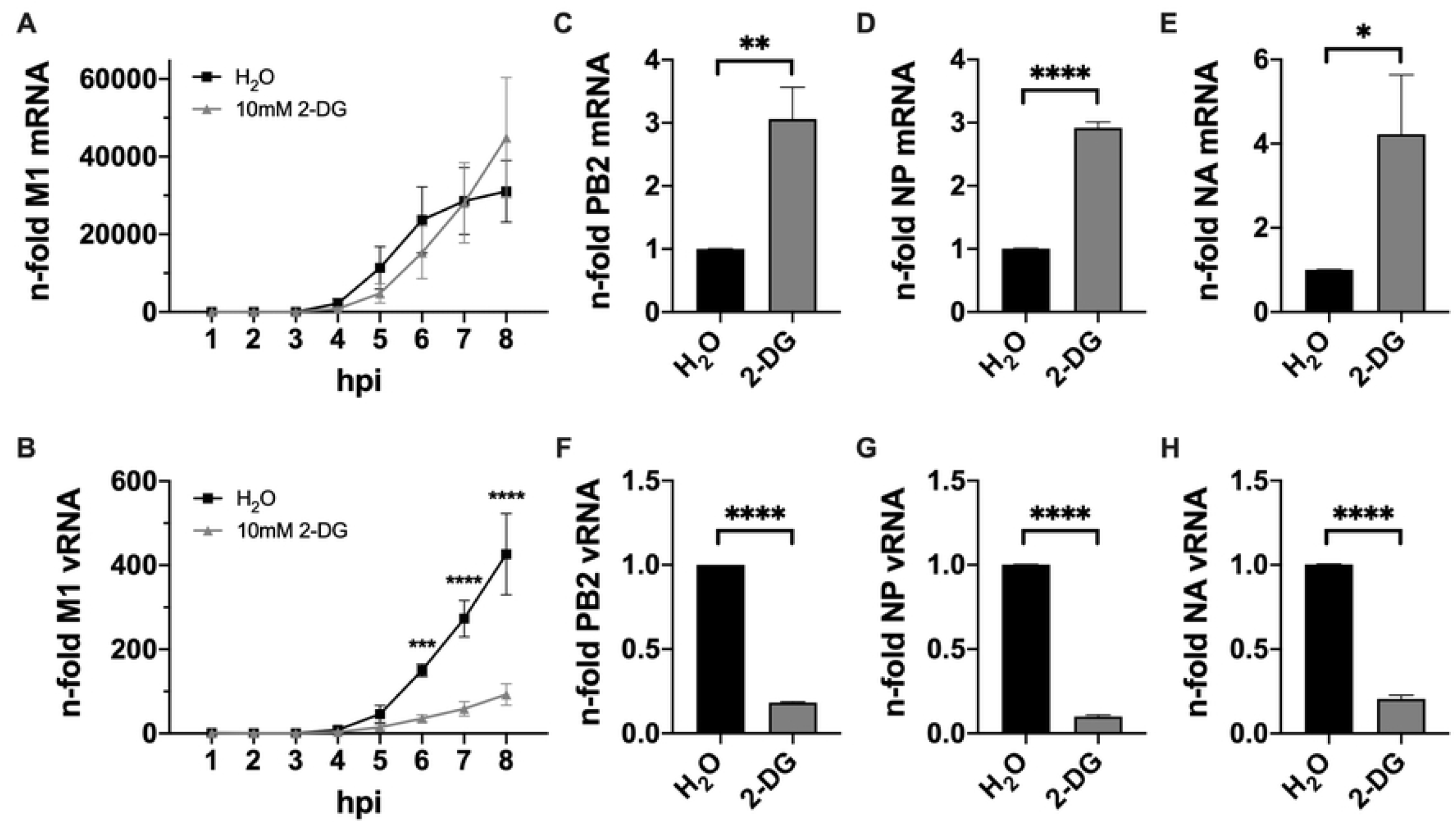
Prolongation of IAV transcription and reduction of replication by 2-DG. 24 h after seeding, A549 cells were infected with SC35M at an MOI of 5 for 30 min and were incubated without or with 10 mM 2-DG in the presence of 25 mM glucose for a total of 8 h. **(A+B)** Each hour or **(C-H)** 8 hpi cells were lysed, their RNA isolated and cDNA synthesized using **(A)** oligo(dT) primers, **(B)** fluA uni12 primers or **(C-H)** specific primers to transcribe mRNA and vRNA of the SC35M gene segments 1 (PB2), 5 (NP) and 6 (NA). Real-time qPCR was performed with two technical replicates per sample. **(A+B)** All values were normalized to the water control 1 hpi or **(C-H)** values of treated samples were normalized to the water control. Depicted are the means ± SD of three independent experiments with three biological replicates per condition and experiment. Statistical significances were determined **(A+B)** via ordinary two-way ANOVA and Sidak’s correction, comparing the treated sample of each time point to its respective water control or **(C-H)** via unpaired t-test. p-values are indicated as follows: < 0.05 = *, < 0.01 = **, < 0.001 = ***, < 0.0001 = ****.

### 2-DG treatment does not affect the replicative capacity of the viral polymerase nor the durability of vRNA

After revealing that reduced vRNA accumulation in the presence of 2-DG was the most crucial consequence of glycolytic interference for viral growth, we wanted to understand this phenomenon more mechanistically. Minigenome systems can be used to explicitly focus on transcription and replication without the dynamic of a full-fledged infection and hence allow to dissect distinct steps of the viral life cycle to a certain degree. Here, minigenome assays were performed as described previously [34] to assess whether 2-DG has a direct influence on the activity of the viral polymerase. For this purpose, we transfected HEK293T cells with plasmids encoding all proteins of the viral ribonucleoprotein (vRNP) complex – PA, PB1, PB2 and NP – together with a reporter plasmid coding for a firefly luciferase under the control of a viral promoter. Another plasmid that constitutively expressed *Renilla* luciferase was co-transfected to serve as a transfection control. Subsequently, those cells were mock-treated or treated with 2-DG and analyzed via luciferase assay. By transfecting two different expression plasmids of the firefly reporter luciferase either vRNA-like or cRNA-like RNA templates were synthesized, which were converted by the transfected and nascent viral proteins. Thus, we were able to analyze the effect of 2-DG on the transcriptional capacity of the viral polymerase (**Fig 5A**) or a potential effect on the replicational capacity of the polymerase since vRNA first had to be synthesized from the cRNA-like template (**Fig 5B**). We observed that transcription was significantly reduced in the presence of 2-DG (**Fig 5A**) which confirms the previously seen 2-DG-induced lower accumulation rate of viral mRNA in the earlier phase of the 8 h kinetic (**Fig 4A**). On the other hand, there was no significant difference of the luciferase signal between the various samples when the cRNA plasmid was transfected (**Fig 5B**). Even though the interpretation here is less straightforward, since the transcription of vRNA in mRNA was also included in this process, this suggested no reduction of the replicational capacity of the viral polymerase by 2-DG.

**Fig 5:**
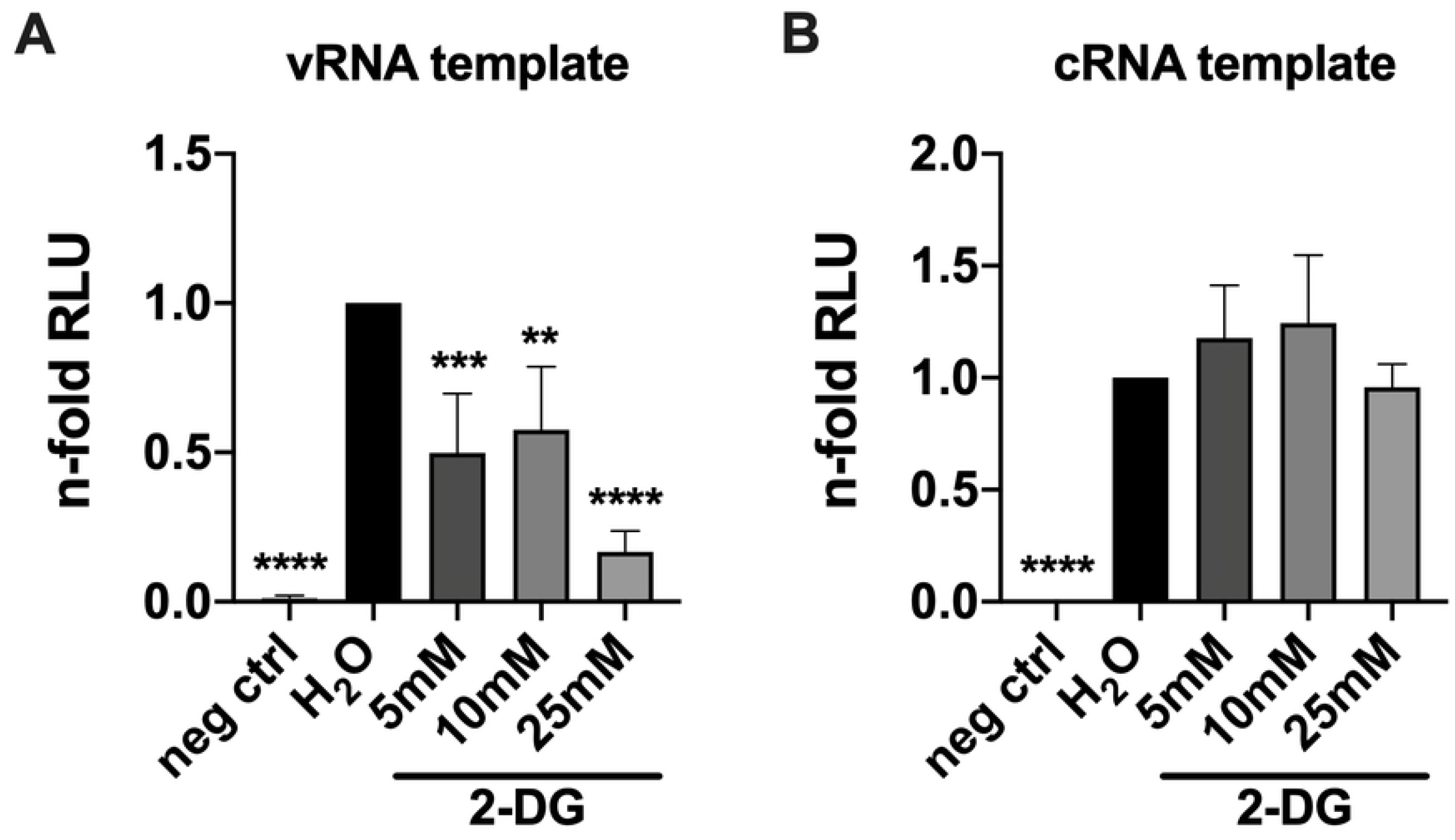
2-DG shows no effect on the replicative capacity of the IAV polymerase. 24 h after seeding, HEK293T cells were transfected with plasmids encoding PA, PB1, PB2 and NP of SC35M, the transfection control *Renilla* luciferase and either a **(A)** vRNA-like or **(B)** cRNA-like template of the *Firefly* luciferase. The negative control was transfected with an empty vector instead of PB2. 4 h later the transfection solution was replaced with medium containing 25 mM glucose and the indicated concentrations of 2-DG for another 20 h. Subsequently, cells were lysed and the n-fold of relative light units (RLU) was measured via luciferase assay and normalized to the water control. Additionally, all values were normalized to their respective transfection control. Statistical significances were determined via unpaired one-way ANOVA and Dunnett’s correction, comparing all other samples to the water control. p-values are indicated as follows: < 0.05 = *, < 0.01 = **, < 0.001 = ***, < 0.0001 = ****.

Additionally, we examined whether the 2-DG treatment potentially affected the durability (e.g., altered stability or rate of degradation) of RNP complexes and performed an assay based on a previous publication [35] in which HEK293T cells were pre-transfected with plasmids encoding all RNP complex proteins. 24 h later they were infected with IAV and subsequently treated with 2-DG and cycloheximide, an inhibitor of translation, for 6 h. This way, the pre-transfected RNP proteins were synthesized and, after IAV infection, formed RNP complexes with the nascent cRNA and vRNA. Strand-specific real-time qPCR revealed that levels of vRNA remained equal between the solvent control and 2-DG-treated samples (**Fig S4**), which indicated no effect of 2-DG on the durability of vRNP complexes.

The data presented in the last two chapters suggested that 2-DG mainly impaired IAV replication and spread by interfering with viral genome replication which was marked by massively reduced levels of vRNA if the inhibitor was applied. However, 2-DG neither had a direct effect on the replicative capacity of the viral polymerase (**Fig 5B**) nor on the durability of vRNP complexes (**Fig S4**).

### IAV infections and glycolytic interference alter the metabolic profile of A549 cells

Given the fact that viral infections affect the cellular metabolism and after revealing that the IAV life cycle is mainly impaired on the level of vRNA synthesis by glycolytic interference, we wanted to get a more comprehensive understanding of metabolic alterations induced by the virus and by a treatment with 2-DG. As we know from the literature [7-9], an IAV infection has profound impacts on the host’s metabolism which especially applies to the glucose metabolism. Since IAV upregulates the glucose metabolism and 2-DG inhibits glycolysis, we expected a (partial) reversion of virus-induced metabolic changes through the inhibitor. Moreover, we were interested in metabolic changes aside from glycolysis. Via hydrophilic interaction liquid chromatography (HILIC) coupled to tandem mass spectrometry (MS/MS), as described previously [36], we analyzed major alterations of the metabolic profile of A549 cells, induced by IAV infection and/or the treatment with 2-DG after 24 h (**Fig 6**).

**Fig 6:**
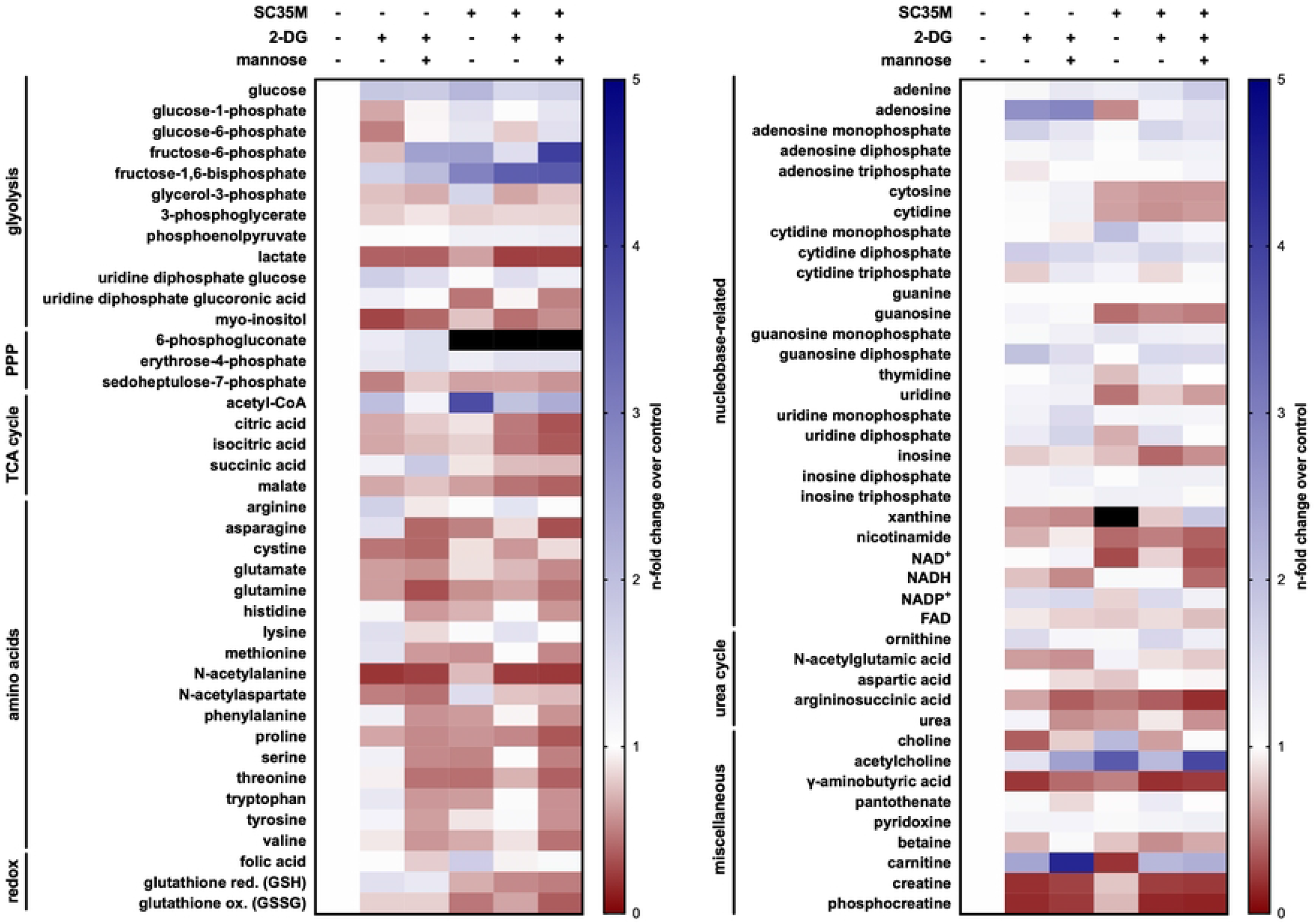
Metabolic alterations induced by IAV infection and glycolytic treatment. A549 cells were mock-infected or infected with SC35M at an MOI of 0.001 and were subsequently incubated in DMEM (containing 25 mM glucose) with or without 10 mM 2-DG and 1 mM mannose as indicated. 24 hpi metabolic activity was quenched and intracellular metabolites were relatively quantified via HILIC-MS/MS. All values have been normalized to the uninfected and untreated control (left column). Darker shades of blue indicate a higher and darker shades of red indicate a lower n-fold of the respective metabolite compared to the control. Black indicates increases higher than 5-fold compared to the control. Depicted are the means of three independent experiments with three biological replicates per condition and experiment. Statistical significances were determined via ordinary two-way ANOVA and Dunnett’s correction, comparing all samples to their respective uninfected and untreated control. The n-folds and p-values are presented in **Table S1**.

In accordance with the literature [7, 8, 10], the levels of glucose and most detected glycolysis intermediates were increased in infected cells, pointing towards an increase of the uptake of glucose and the rate of glycolytic activity. When 2-DG was applied, several glycolytic intermediates (e.g., glucose-6-phosphate (G-6-P) or F-6-P) were detected at decreased concentrations in both, infected and uninfected cells. Counterintuitively, the amount of lactate was decreased in infected cells, which may be explained by an increased efflux upon infection [7, 8] or its metabolization into other intermediates. Independent of an infection, the treatment with 2-DG clearly decreased intracellular lactate. Altogether our data confirmed a virus-mediated upregulation of glycolysis as well as its downregulation in the presence of 2-DG. In combination with our previous data this strengthens the position of metabolic inhibitors as effective antivirals by counteracting virus-induced alterations of the host metabolism.

Other metabolic pathways which are closely connected to glycolysis, such as the PPP or the TCA cycle, revealed some fascinating changes induced by 2-DG treatment or an IAV infection. Among all analyzed metabolites 6-phosphogluconate (6-PG) exhibited the strongest increase upon infection (> 8-fold). The supplementation of 2-DG increased 6-PG concentrations in uninfected and infected cells. This suggested a strong redirection of G-6-P towards the PPP which was probably actively induced by the virus or by the inhibition of GPI by 2-DG. It seems that the oxidative branch of the PPP and thus the direct oxidation of glucose is upregulated upon IAV infection. Similar results have been obtained previously in chicken embryo cells [10]. However, the profiles of detectable downstream intermediates of the non-oxidative PPP differed from each other and were therefore difficult to interpret.

Most of the detected TCA cycle intermediates decreased upon inhibition of glycolysis (abolishment of the anaplerotic function of glycolysis) and during infection. The concentration of acetyl coenzyme A (acetyl-CoA), the linking intermediate between glycolysis and the TCA cycle, was increased in uninfected cells in the presence of 2-DG. But the highest increase of acetyl-CoA was found in untreated but infected cells. Apparently, IAV infections promote the production of large quantities of the important coenzyme. Among amino acids we observed, with only a few exceptions (arginine, lysine and *N*-acetylaspartate), that most of them were decreased during infection. But, while 2-DG led to a decrease of approximately half of the analyzed amino acids it also induced a moderate increase of the other amino acids, independent of an infection. Besides, we noticed that ketogenic or partly ketogenic amino acids were barely or not reduced by 2-DG. Ketogenic amino acids can be catabolized into keto bodies (mostly TCA cycle intermediates such as acetyl-CoA, succinyl-CoA, or fumarate). The amino acids with most severely reduced concentrations after 2-DG treatment all belonged to the group of glucogenic amino acids, which means they can be catabolized into glucose through gluconeogenesis. In favor of this, we also found slightly increased concentrations of pyridoxine (vitamin B6), which is a co-factor for transaminase reactions which convert amino acids into substrates for gluconeogenesis [37, 38]. The inhibition of glycolysis by 2-DG feigned the deprivation of glucose and hence mimicked starvation. Probably this triggered cells to catabolize more glucogenic amino acids. Furthermore, we observed a disturbance of the glutathione equilibrium, one of the most important antioxidant factors for cellular redox homeostasis. In line with this finding, the disruption of glutathione and consequentially the redox homeostasis, as an important factor for IAV pathogenicity, was described before [39-41].

The effect of an IAV infection and especially of 2-DG on many nucleobase-related metabolites (e.g., nucleobases, nucleosides and coenzymes with related structures) was rather mild. Despite the virus-mediated increase in glycolysis, just like Ritter *et al*. reported [8], we observed no significant alteration of ATP levels 24 hpi. Even though to a mild extent, the treatment with 2-DG had the expected effect on intracellular ATP in uninfected cells: 2-DG led to an ATP decrease via inhibition of glycolysis (which even consumes ATP upstream of the inhibition of GPI through the ATP-driven phosphorylation of 2-DG to 2-DG-6-P). The increase of adenosine monophosphate (AMP) in the presence of 2-DG is supported by previous publications reporting of the activation of AMP-activated protein kinase (AMPK) after glycolytic inhibition, which is triggered by a low ATP/AMP ratio [22]. One of the most striking increases upon infection was the 6-fold increase of xanthine which was abolished if 2-DG was administered to infected cells. Despite the decrease of oxidized glutathione (GSSG), the virus-induced increase of xanthine might be linked to the generation of reactive oxygen species (ROS, in this case superoxide) which is an important factor for IAV pathogenicity and proliferation [42]. The catabolism of adenosine generates, among others, xanthine which in turn is a substrate for xanthine oxidase to generate superoxide [43]. In this light, a high catabolic rate of adenosine to generate xanthine during IAV infections may explain the low and high concentration of adenosine and xanthine, respectively, in infected but untreated cells.

Among miscellaneous metabolites we found three candidates which were distinctly affected. The first one is carnitine which is important for the mitochondrial shuttle of fatty acids for β-oxidation and thus the lipid metabolism. Interestingly, in infected but untreated cells carnitine was heavily decreased, suggesting a potential alteration of the lipid metabolism, which was earlier demonstrated in IBV-infected mice [44]. By inhibiting β-oxidation, the virus increases the pool of lipids which can be utilized for the viral envelope, biosynthesis and transport purposes. The other two striking metabolites were creatine and phosphocreatine which were moderately reduced upon IAV infection but heavily reduced in the presence of 2-DG. A main task of these molecules is the conversion of ADP into ATP to sustain energy levels. The strong downregulation of creatine and phosphocreatine might have correlated with the conspicuously mild impact of IAV and 2-DG on ATP concentrations by depleting creatine/phosphocreatine pools in order to maintain sufficient ATP levels.

All described measurements so far aimed to better understand IAV and 2-DG-induced metabolic alterations. However, beside these effects, we also analyzed samples which were additionally supplied with mannose, a C2 epimer of glucose. Since mannose can be converted into fructose-6-phosphate (F-6-P) it should be able to bypass the inhibition by 2-DG to refuel glycolysis. Hence, we expected mannose to reverse some 2-DG-induced effects. Importantly, we observed this reversion, sometimes even followed by an increase, for several glycolytic intermediates (e.g., G-1-P, G-6-P and F-6-P) which demonstrated the antagonistic effect of mannose against glycolytic inhibition by 2-DG. The reduction of ATP in uninfected cells was reversed by mannose, too. However, mannose did not always reverse up-/downregulations of metabolites triggered by 2-DG. For instance, mannose could not or barely reverse the 2-DG-induced decreases of TCA cycle intermediates or alterations among PPP metabolites. Altogether, it seemed that the most pronounced reversions of 2-DG-mediated alterations on the metabolism by mannose took place among intermediates of the glucose metabolism and certain amino acids. Nevertheless, the supplementation of mannose sometimes affected metabolites apart from reversing 2-DG-mediated alterations.

Taken together, these data showed how diversely metabolic pathways are modified during IAV infections and that even metabolites from the same pathway may be affected in different manners. Furthermore, the complex connectivity between pathways or single metabolites became obvious once again. In the context of IAV infections it additionally suggested the potential of glycolytic interference to counteract IAV-induced metabolic changes as well as a function for mannose to regulate 2-DG-mediated effects.

### Mannose circumvents the virus-restricting effect of 2-DG by refueling glycolysis

As described before, glycolysis is closely linked to various other metabolic pathways and its level of activity, as seen in **Fig 6**, can have a strong impact on the abundance of other metabolites. As shown in **Fig 7A** a very close connection exists to the mannose metabolism since F-6-P from glycolysis and mannose-6-phosphate can be converted into each other by the enzyme mannose-6-phosphate isomerase (MPI). Therefore, glucose and mannose should be able to substitute each other for many of their purposes inside a cell, which would also explain some results of the metabolomic data (**Fig 6**). Indeed, the vast majority of mannose is usually shunted to glycolysis to be catabolized. The remaining mannose is mainly utilized for *N*-linked glycosylation [45]. Due to the close connection of glycolysis and *N*-linked glycosylation and since others reported that the antiviral effect of 2-DG originated from the impairment of glycosylation [25, 46] rather than glycolytic inhibition, we aimed to dissect the interplay of these two hexoses in the context of IAV infections and the virus-restricting effects of 2-DG. Since previous publications have shown that 2-DG reduced IAV glycoprotein synthesis [24, 47] and that in general the inhibition of glycosylation by 2-DG could be reversed by low doses of mannose [16, 48], we supplied 2-DG-treated cells with mannose to see if this would reverse the inhibition of viral growth in our cell culture model as well (**Fig S5A**). Indeed, low concentrations of mannose restored viral titers almost completely. We observed this abolishment of the inhibitory function of 2-DG until a 1:10 ratio between mannose (1 mM) and 2-DG (10 mM). To elucidate if the reversal of inhibition can be attributed to mannose being catabolized via glycolysis or being utilized for *N*-linked glycosylation we used the MPI inhibitor MLS0315771 (MLS) to disrupt the link between these two pathways [49]. First, we determined a safe dosage of the inhibitor including potential effects on cell growth, glycolysis, and the formation of infective viral particles. We observed no significant effect on cell proliferation and cell viability but an increase of lactate in the medium in the presence of 50 μM MLS, indicating the safe use of the indicated concentrations and a higher glycolytic rate when the inhibitor is applied (**Fig S5B-D**). The latter can be explained by the fact that MLS prevents the redirection of F-6-P to *N*-linked glycosylation. Therefore, more glucose will be catabolized into lactate via glycolysis. Besides, we observed no significant effect on the production of viral particles (**Fig S5E**). Subsequently, we applied MLS to infected cells which were also treated with 2-DG and mannose (**Fig 7B**). We saw the typical reduction of viral titers when 2-DG alone was applied and the restoration of titers via the addition of mannose. Increasing concentrations of MLS decreased viral titers back to the level of 2-DG-treated samples which suggested that mannose restored IAV propagation mainly by driving glycolysis and not *N*-linked glycosylation. Furthermore, it also confirmed that the inhibition of glycolysis was indeed the primary antiviral mode of action of 2-DG. This got substantiated by the fact that the addition of pyruvate, the final product of glycolysis under physiological conditions, partially restored viral titers after inhibition by 2-DG (**Fig S5F**). To finally confirm the concept of the glycolytic rate as a determinant of IAV replication, we examined the effects of 2-DG, mannose, and MLS on the RNA levels of IAV after a single replication cycle of 8 h. The pattern of M1 vRNA accumulation (**Fig 7C**) strikingly resembled the pattern of viral titers (**Fig 7B**). The treatment with 2-DG led to a highly significant reduction of vRNA which was almost completely restored to the control value by supplementation of mannose. The additional administration of MLS, however, decreased the vRNA value to a similar extent as 2-DG alone did. Regarding viral mRNA accumulation (**Fig 7D**) we observed the typical slight increase after treatment with 2-DG, but barely a return to the control value when mannose was added as well. This only happened when also MLS was supplemented.

**Fig 7:**
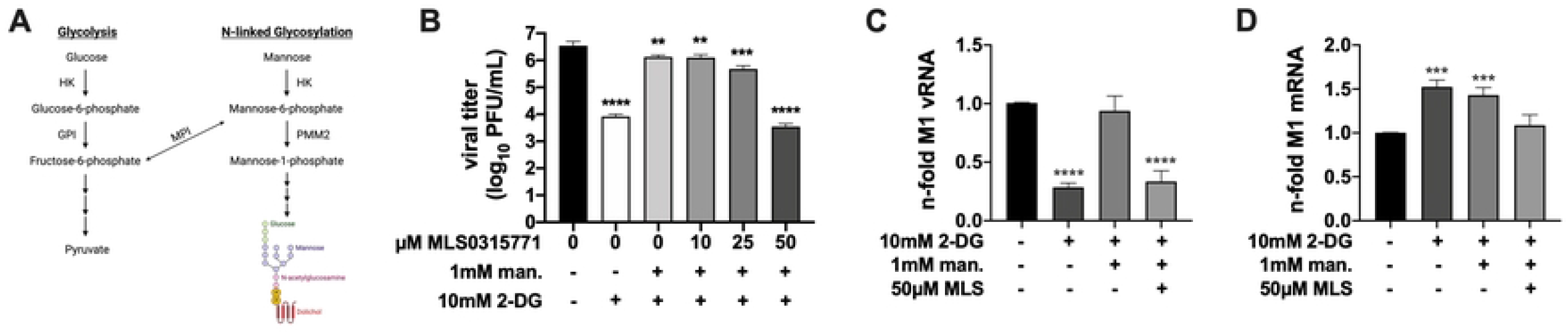
Mannose counteracts 2-DG by refueling glycolysis. **(A)** The metabolic pathways of glycolysis and *N*-linked glycosylation are closely connected via mannose-6-phosphate isomerase (MPI). Other enzymes depicted here are hexokinase (HK), glucose-6-phosphate isomerase (GPI), and phosphomannomutase 2 (PMM2). **(B-D)** 24 h after seeding, A549 cells were infected with SC35M at an MOI of **(B)** 0.001 or **(C+D)** 5 for 30 min and were incubated with 25 mM glucose and the indicated concentrations of 2-DG, mannose, and the mannose-6 phosphate isomerase inhibitor MLS0315771 (MLS) for a total of **(B)** 24 h or **(C+D)** 8 h. Subsequently, **(B)** supernatants were collected to determine viral titers via plaque assay or **(C+D)** cells were lysed, their RNA isolated and cDNA synthesized using either **(C)** fluA uni12 primers to transcribe vRNA or **(D)** oligo(dT) primers to transcribe mRNA. Real-time qPCR was performed with two technical replicates per sample and values of treated samples were normalized to the untreated control. In case of mRNA detection all results were additionally normalized to a GAPDH control. Depicted are the means ± SD of three independent experiments with three biological replicates per condition and experiment. Statistical significances were determined via unpaired one-way ANOVA and Dunnett’s correction, comparing all treated samples to the untreated control. p-values are indicated as follows: < 0.05 = *, < 0.01 = **, < 0.001 = ***, < 0.0001 = ****.

Summarizing, these data corroborated that the antiviral activity of 2-DG mainly derived from a strong impairment of the synthesis of viral genomic RNA by reducing the glycolytic rate of infected host cells. Moreover, by directly or indirectly inhibiting or fueling glycolysis we found a way to turn viral reproduction on and off to a certain degree.

## 3. Discussion

Understanding the diverse interplay between the host cell metabolism and viral intruders is of importance since it may create potential new strategies to counteract viral infections. In our study we were able to improve our comprehension of metabolic virus-host interactions as well as the mode of action of glycolytic interference on the life cycle of IAV. We observed profound changes of the whole metabolic profile of infected cells (**Fig 6**), including especially upregulated amounts of many intermediates of glycolysis. By applying 2-DG, a potent inhibitor of glycolysis, many virus-induced metabolic alterations could be reversed which indicated the inhibitor’s counteraction against viral manipulations of the host. Furthermore, we showed the severe impact of glycolytic interference by 2-DG on the propagation of IAV *in vitro* (**Fig 1A+S1A**). The reduction of virus titers reached up to 4.5 orders of magnitude and hence was similar or even exceeded the effectivity of other antiviral compounds [50, 51].

During our search for the point of interference within the viral life cycle we deduced that viral protein synthesis played a subordinate role, because protein accumulation was rather mildly affected within one replication cycle (**Fig 2B**). Interestingly, we did not at all observe a decrease in cellular protein accumulation after a 2-DG treatment, shown by steady signals of the cellular control protein in our western blots and via *Renilla* luciferase reporter assay (**Fig 2**). This may be indicative of a more selective effect which rather applies to viral than cellular protein translation. According to the data of **Fig 4** and **5** we assume that the predominant mechanism which is responsible for the strong reduction of IAV multiplication is a 2-DG-mediated interference with the dynamic regulation that switches the viral polymerase from a transcriptase to a replicase. Even though one hypothesis proposed that viral transcription and replication are stochastic without a switch mechanism [35] many studies suggest the opposite. The switch process of the polymerase is still not fully understood and is probably a multifactorial process determined by several viral and host factors (summarized in [31]). NP seems to be one of the key factors in this context and was shown to have a stimulatory function on viral polymerase activity via a direct interaction with it [52-55]. Additionally, NS1 and nuclear export protein (NEP, also known as NS2) are presumably implicated in viral replication [56-58]. Furthermore, small viral RNAs (svRNAs), which resemble the 5’ end of vRNAs, have been linked to the regulation of viral replication [59-62]. It’s been hypothesized that the role of svRNAs in viral replication is the association with a second and trans-acting polymerase which binds the 5’ end of newly synthesized vRNA [31]. Even though it once was postulated that host factors are not required to initiate viral replication [54], many candidates that can associate with vRNP components [31, 55, 63-66] and thereby potentially influence the process, such as the recently described acidic nuclear phosphoprotein 32 (ANP32) [67-69], have been identified. Since the nuclear matrix and chromatin of infected cells were postulated to constitute a platform for viral transcription and replication [70-72], various potential host factors are associated with these sub-nuclear structures [73-76]. Linking the described regulators of IAV polymerase activity and the here presented data, it is quite possible that metabolic interference via 2-DG impairs the IAV replication-associated function or the expression of one or several of these viral or host factors. After all we know, however, it is also possible that there is no strong switching mechanism controlling viral transcription or replication. Potentially the abundance of both processes is basically stochastic but can be modulated in favor of transcription or replication in a time-dependent manner. Combining the insights from previous publications with our data it is imaginable that the antiviral effect of 2-DG operates in several steps. One scenario could be that inhibition by 2-DG leads to a primary antiviral effect by interfering with the function of the initial transcription and replication complexes which could explain the generalized lower levels of mRNA and vRNA until 7 hpi (**Fig 4**). A secondary effect could be the seemingly selective impairment of the synthesis of some viral proteins. This includes at least PA of the polymerase or RNP complex (**Fig 2B**). Consequently, a lack of nascent polymerase complexes may have a stronger impact on replication than transcription since replication requires a second polymerase for the binding of nascent cRNA and vRNA strands. Alternatively or additionally, treatment with 2-DG might impair the synthesis of any of the afore-mentioned modulators of the viral polymerase which may contribute to the prolonged phase of transcription and the clear reduction of replication. Of course, the variety of potential 2-DG-mediated influences on viral replication is huge and on top of that we cannot fully exclude an off-target interaction which may play a role here. However, the latter seems highly unlikely based on the data we generated through the supplementation of mannose and MLS in the presence of 2-DG (**Fig 7**). It will be interesting to examine if and how severely 2-DG influences the expression or interactions of the afore-mentioned viral and cellular factors with the complex replication machinery of IAV.

Furthermore, our data suggest that the predominant antiviral mode of action of 2-DG is the inhibition of glycolysis. Decades ago it has been postulated that the impairment of *N*-linked glycosylation is responsible for the antiviral effect of 2-DG [46]. The fact that inhibition of the enzyme MPI, which links glycolysis and glycosylation, abolished the restoration of viral titers and vRNA levels by mannose after treatment with 2-DG (**Fig 7B+C**) lets us oppose this view. Our data indicate that the positive effect of mannose on IAV replication mainly (but not necessarily exclusively) derives from fueling glycolysis via its conversion into F-6-P by MPI. The partial recovery of the ATP/AMP ratio back to the physiological level in the presence of 2-DG and mannose (**Fig 6**) supports this theory. Moreover, the partial restoration of viral titers by the supplementation of pyruvate after inhibiting glycolysis substantiates the assumption that glycolysis and its intermediates are crucial for virus reproduction. Probably the availability of glycolytic intermediates, which are needed to fuel other pathways and to synthesize macromolecules such as nucleotides and amino acids, is the most critical factor. Extrapolations predicted only a very minor extra demand for energy (∼1% of the total energetic budget of a eukaryotic cell) to synthesize viral progeny during the characteristic time of an influenza infection [77]. Therefore, we assume that a potential role of ATP in viral replication may rather not be its availability for synthesis reactions.

As reviewed previously [78], 2-DG has various direct and indirect mechanism by which it can negatively affect normal cellular functions (e.g., inhibition of glycolysis and glycosylation or induction of AMPK and UPR). Therefore, a certain cytotoxicity – which heavily depends on the dose and type of administration as well as the type of cell, tissue, or organism – must be considered. However, we could demonstrate the tolerability and the quickness of effectivity of the antimetabolite in immortalized and primary cells (**Fig 1C-F, S1B, S2B+C**). Our *in vitro* data and previous reports [26, 27] support the performance of more *in vivo* studies and clinical trials to assess the safety of 2-DG and its efficiency to treat virus infections in model organisms or even humans. Several such studies have already reported the safety of 2-DG in animal models in the context of other virus infections [79] or different fields of research [80-82], especially when administered in continuous low doses. This could even be confirmed in clinical trials [83, 84]. Very recent phase II and III clinical trials in India [85, 86] demonstrated the safety and effectiveness of 2-DG when applied in addition to the standard of care to treat severe COVID-19 patients. As studies in which a virus infection was more successfully treated in humans through metabolic interference, these clinical trials may become a milestone in the development of host-targeted metabolic drugs as antivirals. However, some studies [87, 88] and its poor pharmacokinetic properties, e.g., its short plasma half-life [89], suggest that 2-DG itself may never become a licensed drug. Nevertheless, it is a useful tool to examine the principles of glycolytic interference and novel 2-DG analogs or other glycolytic inhibitors possibly boast a better pharmacological suitability [90]. Since dependence on the host metabolism is a universal feature of all viruses, differential and strictly determined metabolic treatments may be able to alleviate all types of virus infections in the future. However, before this may become reality, we need to gain a more comprehensive understanding of metabolism-related virus-host interactions, including virus-induced metabolic modifications, specific metabolic needs of different viruses and how exactly metabolic treatments affect the viral life cycle as well as the host. We are positive that this specific field of research deserves more attention to elaborate metabolic interference and make it become a realistic and sensible treatment option in the future.

## 4. Experimental procedures

### 4.1 Cell lines and viruses

Human adenocarcinomic alveolar basal epithelial cells (A549, American type culture collection (ATCC®), CCL-185™) and human embryonic kidney (HEK) 293t cells (ATCC®, CRL-3216™) were cultured in the high glucose variant of Dulbecco’s modified Eagle’s medium (DMEM, Sigma-Aldrich, D5796) supplemented with 10 % fetal bovine serum (FBS). Madin-Darby canine kidney (MDCK) II cells (Institute of Virology, WWU Muenster, Germany) were cultured in minimum essential medium (MEM, Sigma-Aldrich, M4655) supplemented with 10 % fetal bovine serum (FBS). The primary cells human bronchial epithelial cell (HBEpC, PromoCell, C-12640) were cultured in airway epithelial cell growth medium (AECGM, PromoCell, C-21060). Tumor-free human lung explants were obtained from various donors right after surgery at the University Hospital Muenster and were cultured in Roswell Park Memorial Institute-1640 medium (RPMI-1640, Sigma-Aldrich, R8758) supplemented with 100 U/mL penicillin and 0.1 mg/mL streptomycin. The donors gave written consent for the tissue to be used for scientific purposes. Ethical approval was given by the Deutsche Ärztekammer (AZ: 2016-265-f-S). All cells were kept at 37 °C and 5 % CO_2_. Mouse-adapted A/Seal/Massachusetts/1/80 H7N7 (SC35M) and A/Panama/2007/1999 H3N2 (Pan/99) are recombinant influenza A virus (IAV) strains which were propagated in MDCK II cells.

### 4.2 Infection and treatment

Viruses were diluted to the desired multiplicity of infection (MOI) in phosphate-buffered saline (PBS) supplemented with 0.2 % bovine serum albumin (BSA), 1 mM MgCl_2_, 0.9 mM CaCl_2_, 100 U/mL penicillin and 0.1 mg/mL streptomycin. Cells were washed once with PBS and incubated for 30 min at 37 °C and 5 % CO_2_ with the respective amount of virus. Afterwards A549 and HEK293T cells were washed once more with PBS and then incubated for the depicted periods in DMEM (Thermo Fisher Scientific, A14430) containing 0.2 % bovine serum albumin (BSA), 100 U/mL penicillin and 0.1 mg/mL streptomycin, 25 mM D-glucose, 2 mM L-glutamine and the respective concentration of inhibitor/supplement. The medium did not contain sodium pyruvate, HEPES and phenol red. HBEpCs were washed once with PBS after an infection and incubated in AECGM, containing the respective amounts of inhibitor/supplement for the depicted periods of the experiments. Human lung explants (∼100 mg) were infected with 2 × 10^5^ infectious virus particles as described previously [91], but without any interferon or bafilomycin. After washing the tissue 1 hpi, it was incubated in fresh RPMI supplemented with 2 mM L-glutamine, 100 U/mL penicillin, 0.1 mg/mL streptomycin, 0.1 % bovine serum albumin and the indicated concentrations of inhibitor. 2-deoxy-D-glucose (2-DG, Sigma-Aldrich, D8375), D-(+)-mannose (Sigma-Aldrich, M6020) and sodium pyruvate (Sigma-Aldrich, P5280) were dissolved in H_2_O to 1 M (2-DG and mannose) and 2 M (sodium pyruvate) stock solutions. MLS0315771 (MedChemExpress, HY-112945) was dissolved in dimethyl sulfoxide (DMSO) to a stock concentration of 10 mM. For the stimulation of immune responses via RNA transfection, RNA was isolated from mock-infected and SC35M-infected (MOI of 5) cells 8 hpi, as described in **4.7**. 100 ng RNA per well was transfected using HiPerFect Transfection Reagent (QIAGEN) according to the manufacturer’s protocol for 6 h in the presence of the depicted inhibitor concentrations.

### 4.3 Plaque titration

After the indicated periods of infection, the supernatants were collected and used to determine the number of infectious virus particles. Confluent MDCK II cells were infected with serial dilutions of the supernatants in PBS containing 0.2 % bovine serum albumin (BSA), 1 mM MgCl_2_, 0.9 mM CaCl_2_, 100 U/mL penicillin and 0.1 mg/mL streptomycin for 30 min at 37 °C and 5 % CO_2_. Subsequently the supernatants were replaced with MEM/BA containing 0.21 % BSA, 0.21 % NaHCO_3_, 1 mM MgCl_2_, 0.01 % DEAE-dextran, 0.9 mM CaCl_2_, 100 U/ml penicillin, 0.1 mg/ml streptomycin and 0.9 % purified agar. After an incubation for 2-3 days at 37 °C and 5 % CO_2_ the overlay was removed and cells were stained with a Coomassie staining solution (45 % ddH_2_O (v/v), 45 % methanol (v/v), 10 % acetic acid (v/v) and 0.25 % Coomassie Brilliant blue R-250 (w/v)). Cell free plaques in the monolayer were counted as plaque-forming units per milliliter (PFU/mL).

### 4.4 Cytotoxicity assays

Potential cytotoxic effects of inhibitors were assessed by three different methods: lactate dehydrogenase (LDH) assay, trypan blue staining and flow cytometry. LDH assays were performed with the CytoSelect LDH cytotoxicity assay kit (Bio Cat, CBA-241-CB) according to the manufacturer’s manual. Trypan blue exclusion was done by mixing a 0.4 % trypan blue dye (Invitrogen) 1:1 with a sample’s cell suspension and having the automated cell counting machine Countess™ II (Invitrogen) determine the number of living cells. Determination of living cells via flow cytometry is described below in section **4.10**.

### 4.5 Glycolytic rate test

The induced assay version of the glycolytic rate test (Agilent, Kit 103344-100) was performed with a Seahorse XFe96 Analyzer (Agilent) according to the manufacturer’s instructions. The assay medium was supplemented with 25 mM D-glucose and 2 mM L-glutamine to match other experimental conditions. Concomitantly, the final injection of 2-DG was set to 125 mM. After three measured points to obtain the basal glycolytic level, the indicated concentrations of inhibitor were injected and the glycolytic rate was measured for 1 h before continuing with the standard procedure.

### 4.6 Lactate assay

To determine the concentration of lactate in the supernatants of samples and thus have an indirect assay to assess glycolytic activity, the L-Lactate Assay Kit II (PK-CA577-K607) from PromoCell was used according to the manufacturer’s instruction.

### 4.7 Reverse transcription and quantitative real-time PCR

At the end of an infection and/or treatment period, total RNA was isolated using the RNeasy® Plus Mini Kit (Qiagen). The procedure was done according to the manufacturer’s manual. Reverse transcription was performed with the RevertAid™ H Minus Reverse Transcriptase (Thermo Fisher Scientific) and oligo(dT) primers (Eurofins Genomics) for detection of mRNA or a fluA uni12 forward primer [92] (Sigma-Aldrich, 5’-AGCAAAAGCAGG-3’) to detect vRNA according to the manufacturer’s protocol. The obtained cDNA was used for real-time qPCR with a LightCycler® 480 II (Roche) and Brilliant III SYBR® Green (Agilent) according to the manufacturer’s instructions. The following primers were used during qPCR: influenza matrix protein M1 forward (5’-AGA TGA GTC TTC TAA CCG AGG TCG-3’) and reverse (5’-TGC AAA AAC ATC TTC AAG TCT CTG-3’), IL-6 forward (5’-AGA GGC ACT GGC AGA AAA CAA C-3’) and reverse (5’-AGG CAA GTC TCC TCA TTG AAT CC-3’), CXCL8 forward (5’-ACT GAG AGT GAT TGA GAG TGG AC-3’) and reverse (5’-AAC CCT CTG CAC CCA GTT TTC-3’), DDX58 forward (5’-CCT ACC TAC ATC CTG AGC TAC AT-3’) and reverse (5’-TCT AGG GCA TCC AAA AAG CCA-3’), MxA forward (5’-GTT TCC GAA GTG GAC ATC GCA-3’) and reverse (5’-GAA GGG CAA CTC CTG ACA GT-3’) and human glyceraldehyde 3-phosphate dehydrogenase (GAPDH) forward (5’-GCA AAT TCC ATG GCA CCG T-3’) and reverse (5’-GCC CCA CTT GAT TTT GGA GG-3’). GAPDH, as a housekeeping gene, was used for the normalization of PCR results. The relative n-fold was calculated using the 2^-ββCT^ method [93].

### 4.8 Strand-specific quantitative real-time RT-PCR

Total RNA was isolate as described in **4.7**. Reverse transcription was performed by using Maxima™ Reverse Transcriptase (Thermo Fisher Scientific) according to the manufacturer’s instructions and specific primers (Eurofins Genomics) for the different types of RNA as reported previously [33]. The following primers were designed according to the SC35M sequences DQ266097, DQ226096 and DQ266095 (Influenza Research Database) and used for cDNA synthesis and qPCR of the SC35M segments 1, 5 and 6:

**Table 1:**
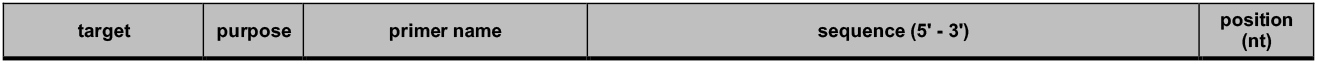
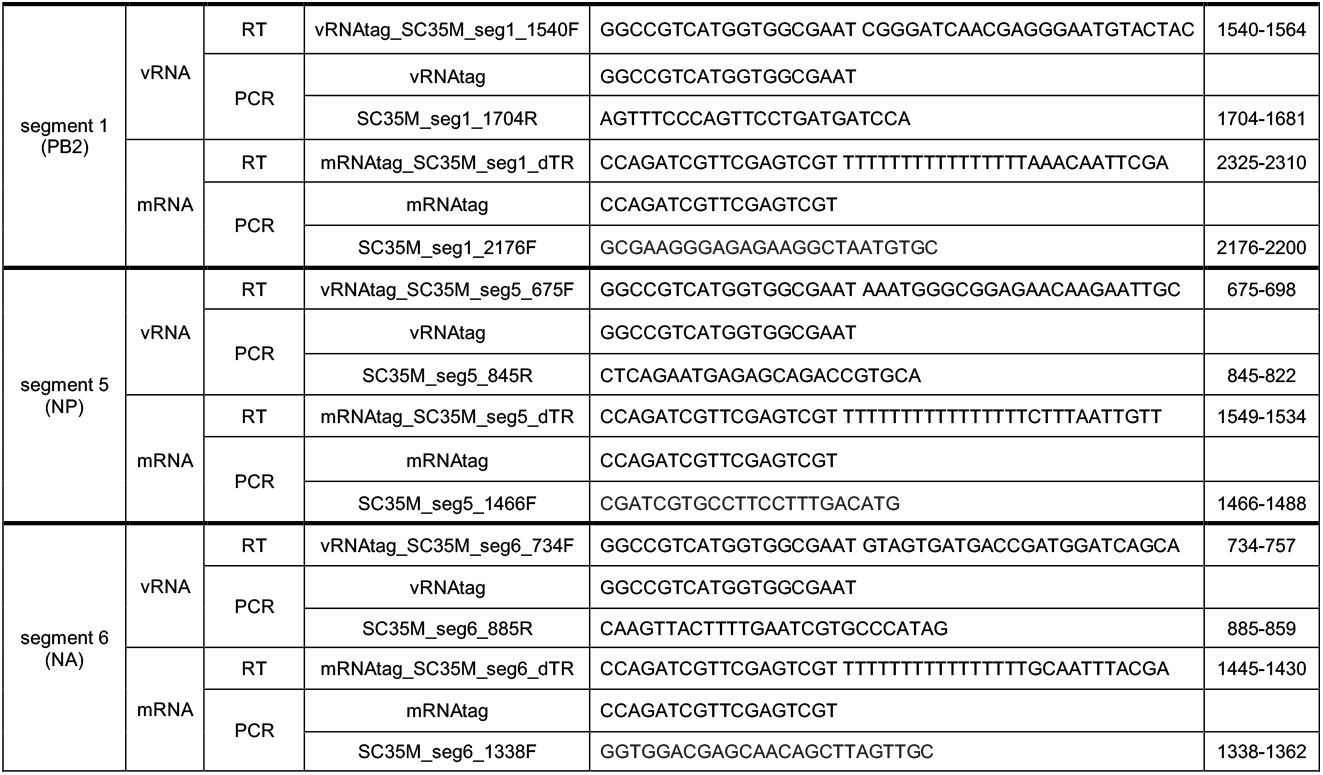
Primer for strand-specific real-time qPCR subdivided into their use in cDNA synthesis via reverse transcription and PCR.

### 4.9 Western blot

Samples were lysed at 4 °C with radioimmunoprecipitation assay (RIPA) buffer (25 mM Tris-HCl pH 8, 137 mM NaCl, 10 % glycerol, 0.1 % SDS, 0.5 % NaDOC, 1 % NP-40, 2 mM EDTA pH 8, 200 μM Pefabloc®, 5 μg/mL aprotinin, 5 μg/mL leupeptin, 1 mM sodium orthovanadate and 5 mM benzamidine). Cell debris was removed via centrifugation and protein concentrations were determined by Bradford assay. Samples were adjusted to the same protein concentration, mixed with the appropriate amount of Laemmli sample buffer and then proteins were separated and visualized by sodium dodecyl sulfate polyacrylamide gel electrophoresis (SDS-PAGE) and western blot analysis. The following primary antibodies were used to detect their respective proteins: ERK2 (rabbit, polyclonal, Santa Cruz, sc-154), M1 (mouse, monoclonal, Biorad, MCA401), NP (rabbit, polyclonal, GeneTex, GTX125989), NS1 (rabbit, polyclonal, GeneTex, GTX125990), and PA (rabbit, polyclonal, GeneTex, GTX125932). ERK2 served as the loading control for whole cell lysates. Fluorescence signals were visualized by using fluorophore-labelled secondary antibodies: IRDye® 680RD Donkey anti-Mouse (LI-COR, 926-68072), IRDye® 680RD Donkey anti-Rabbit (LI-COR, 926-68073), IRDye® 800CW Donkey anti-Mouse (LI-COR, 926-32212), and IRDye® 800CW Donkey anti-Rabbit (LI-COR, 926-32213). Images were taken with the ODYSSEY® F_C_ Imaging System (LI-COR).

### 4.10 Flow Cytometry

At the end of an infection with or without treatment, cells were trypsinized and subsequently stained for analysis via flow cytometry with the FACSCalibur (Becton Dickinson) flow cytometer. At first, cells were stained with eBioscience™ Fixable Viability Dye eFluor™ 660 (Invitrogen, 65-0866-14) for 30 min at 4 °C in the dark. Afterwards the samples were fixated and permeabilized for 20 min and 60 min at 4 °C in the dark using BD Cytofix/Cytoperm™ solution and BD Perm/Wash™ solution (BD Biosciences), respectively. Intracellular staining of influenza A nucleoprotein was done by applying the anti influenza A (nucleoprotein) – FITC antibody (OriGene, AM00924FC-N) for 60 min at 4 °C in the dark. FlowJo software v10 (Becton Dickinson) was used to analyze the data obtained by flow cytometry. 10^5^ cells of each sample were analyzed. The gating strategy is displayed in **Fig S6**.

### 4.11 Minigenome assay

Using Lipofectamine™ 2000 (Invitrogen), HEK293T cells were transfected with polymerase II-driven plasmids coding for PA, PB1, PB2, NP of SC35M as well as the transfection control *Renilla* luciferase. A sixth plasmid was one of two polymerase I-driven plasmids encoding either a vRNA-like or cRNA-like firefly luciferase template. 4 h post transfection the medium was replaced with DMEM (Thermo Fisher Scientific, A14430) containing 0.2 % BSA, 100 U/mL penicillin and 0.1 mg/mL streptomycin, 25 mM D-glucose, 2 mM L-glutamine and the respective concentration of 2-DG. 24 h post transfection the Dual-Luciferase® Reporter Assay System (Promega) was used according to the manufacturer’s manual. For measurements of relative light units (RLU) the luminometer MicroLumat*Plus* LB 96V (Berthold Technologies) and the software WinGlow (Berthold Technologies) were used.

### 4.12 RNP durability assay

HEK293T cells were transfected with plasmids coding for PA, PB1, PB2 and NP of SC35M using Lipofectamine™ 2000 (Invitrogen). 24 h post transfection cells were infected with SC35M at an MOI of 5 (see **4.2**) and incubated with or without cycloheximide (100 μg/mL) and various concentrations of 2-DG. 6 hpi cell lysates were taken and subjected to strand-specific quantitative real-time RT-PCR (see **4.8**).

### 4.13 Metabolic profiling by HILIC-MS/MS

24 h after seeding 1.5 × 10^6^ A549 cells in 6 cm dishes, they were mock-infected or infected with SC35M at an MOI of 0.001. 24 hpi cells were washed twice with PBS and 400 μL pre-cooled (4-8 °C) acetonitrile (ACN)/water (4+1, v/v) including 50 μM D-phenylglycine as internal standard was added for metabolic quenching. Until further preparation the samples were kept at 4-8 °C. Cells were then detached using a sterile cell scraper. The dish was washed with additional 800 μL ACN/water (4+1, v/v) and pooled with the respective cell sample. Further preparation of samples as well as chromatographic and mass spectrometric analysis were performed as described previously [36].

## 5. Acknowledgement

J.K is a member of CiM-IMPRS, the joint graduate school of the Cells in Motion Interfaculty Centre, University of Muenster, Germany and the International Max Planck Research School-Molecular Biomedicine, Muenster, Germany. Besides, we acknowledge technical support from L. Schürmann.

---

**Fig S1: Effects of 2-DG on IAV propagation, cell growth and immune induction**. 24 h after seeding, A549 cells were infected with SC35M at an MOI of **(A+B)** 0.01 or **(C+D)** 0.001 for 30 min and incubated in the presence of the indicated concentrations of 2-DG or its solvent water for 24 h. Subsequently, cells were **(A+B)** stained with an NP antibody and a live/dead marker and analyzed via flow cytometry or **(C+D)** detached to assess the number of living cells as well as the viability via trypan blue exclusion in an automated cell counter. **(E-H)** Uninfected cells were transfected with cellular or viral RNA and treated with the indicated 2-DG concentrations for 6 h. Subsequently, cells were lysed, their RNA isolated and cDNA synthesized using oligo(dT) primers to transcribe mRNA. Real-time qPCR was performed with two technical replicates per sample and values of all other samples were normalized to the unstimulated water control. Additionally, all results were normalized to a GAPDH control. **(A-H)** Depicted are the means ± SD of three independent experiments with three biological replicates per condition and experiment. Statistical significances were determined via **(A, C, D)** unpaired one-way ANOVA and Dunnett’s correction, comparing all treated samples to the water control or **(B, E-H)** ordinary two-way ANOVA with Dunnett’s correction, comparing all treated samples of both groups to their respective water control. p-values are indicated as follows: < 0.05 = *, < 0.01 = **, < 0.001 = ***, < 0.0001 = ****.

**Fig S2: Effects of 2-DG on human primary cells and IAV propagation. (A)** Human lung explants were infected with 2 × 10^5^ SC35M particles for 30 min. Afterwards they were incubated with 11.1 mM glucose and the indicated concentrations of 2-DG and supernatants were collected 1, 24 and 48 hpi to determine viral titers via plaque assay. **(B-I)** After reaching ≈ 90 % confluency **(B)** uninfected HBEpCs were treated with the indicated concentrations of 2-DG or its solvent water for 24 h. Afterwards the supernatants were used to perform LDH assays to determine the relative cytotoxicity of the treatment. **(C-I)** HBEpCs were infected with SC35M at an MOI of **(C)** 1, **(D, F, H)** 0.01 or **(E, G, I)** 5 for 30 min and were incubated with 6 mM glucose and the indicated concentrations of 2-DG for a total of **(D, F, H)** 24 h or **(E, G, I)** 8 h. Subsequently, **(C-E)** supernatants were used to **(C)** perform lactate assays in order to indirectly assess the glycolytic activity and **(D+E)** determine viral titers via plaque assay. **(F-I)** Additionally, cells were lysed, their RNA isolated and cDNA synthesized using either **(F+H)** oligo(dT) primers to transcribe mRNA or **(H+I)** fluA uni12 primers to transcribe vRNA. Real-time qPCR was performed with two technical replicates per sample and values of treated samples were normalized to the water control. In case of mRNA detection, all results were additionally normalized to a GAPDH control. Depicted are the means ± SD of three independent experiments with three biological replicates per condition and experiment. Statistical significances were determined via **(A-C)** ordinary two-way ANOVA and Dunnett’s correction, comparing each treated sample to its respective water control. **(D-I)** Other significances were determined via unpaired one-way ANOVA and Dunnett’s correction, comparing all treated samples to the water control. p-values are indicated as follows: < 0.05 = *, < 0.01 = **, < 0.001 = ***, < 0.0001 = ****.

**Fig S3: Impairment of IAV replication by 2-DG is not strain-specific**. 24 h after seeding, A549 cells were infected with Pan/99 at the depicted MOIs for 30 min and were incubated with the indicated concentrations of 2-DG or its solvent water for a total of **(A-C)** 24 h or **(D+E)** 8 h. Subsequently, **(A)** supernatants were collected to determine viral titers via plaque assay or **(B-E)** cells were lysed, their RNA isolated and cDNA synthesized using either **(B+D)** oligo(dT) primers to transcribe mRNA or **(C+E)** fluA uni12 primers to transcribe vRNA. Real-time qPCR was performed with two technical replicates per sample and values of treated samples were normalized to the water control. In case of mRNA detection, all results were additionally normalized to a GAPDH control. Depicted are the means ± SD of three independent experiments with three biological replicates per condition and experiment. Statistical significances were determined via unpaired one-way ANOVA and Dunnett’s correction, comparing all treated samples to the water control. p-values are indicated as follows: < 0.05 = *, < 0.01 = **, < 0.001 = ***, < 0.0001 = ****.

**Fig S4: 2-DG does not impair IAV vRNA durability**. 24 h after seeding, HEK293T cells were transfected with plasmids containing the SC35M sequences of PA, PB1, PB2 and NP. 4 h later the transfection solution was replaced with fresh medium for another 20 h. Subsequently, cells were infected with SC35M at an MOI of 5 for 30 min and were incubated with the indicated concentrations of 2-DG and 100 μg/mL cycloheximide. A negative control was previously transfected with an empty vector instead of PA while a positive control was not treated with cycloheximide. 6 hpi, cells were lysed, their RNA isolated and cDNA synthesized using specific primers to transcribe vRNA of the SC35M gene segment 6 (NA). Real-time qPCR was performed with two technical replicates per sample. Statistical significances were determined via unpaired one-way ANOVA and Dunnett’s correction, comparing all other samples to the water control. p-values are indicated as follows: < 0.05 = *, < 0.01 = **, < 0.001 = ***, < 0.0001 = ****.

**Fig S5: Effects of mannose, MLS0315771, and pyruvate on IAV propagation and A549 cells**. 24 h after seeding, A549 cells were infected with SC35M at an MOI of 0.001 or for 30 min and incubated in the presence of the indicated concentrations of metabolites and inhibitors or their solvents for a total of 24 h. Subsequently, **(A, D-F)** supernatants were collected to determine **(A, E, F)** viral titers via plaque assay and **(D)** extracellular lactate concentrations via lactate assay or **(B+C)** cells were detached to assess the number of living cells and the viability via trypan blue exclusion and an automated cell counter. Depicted are the means ± SD of three independent experiments with three biological replicates per condition and experiment. Statistical significances were determined via unpaired one-way ANOVA and Dunnett’s correction, comparing **(B-E)** all treated samples to the DMSO control or **(A+F)** all other samples to the 2-DG-treated sample (white bar). p-values are indicated as follows: < 0.05 = *, < 0.01 = **, < 0.001 = ***, < 0.0001 = ****.

**Fig S6: Gating strategy for the quantification of uninfected versus infected and living versus dead cells**. A549 cells were infected with SC35M at an MOI of 0.01. Directly after the infection, cells were mock-treated or treated with 2-DG. 24 hpi cells were stained with a viability dye and an NP antibody and were quantified via flow cytometry. At first cells were pre-gated according to their FSC/SSC appearance. Then these cells were sub-classified to discriminate between uninfected and infected cells as well as living and dead cells. Representative dot plots are depicted to exemplify the gating strategy used for data analysis in Fig S1A+B.

**Table S1: n-fold changes over control and statistical significances of Fig 6**.

